# Aerosol deposition increases conductance to water in immobilized stomata closed with abscisic acid or opened with fusicoccin

**DOI:** 10.1101/2023.12.17.571850

**Authors:** David A. Grantz, Chia-JuEllen Chi, Juergen Burkhardt

## Abstract

Hypothesized effects of aerosol deposition on plant water balance have been difficult to establish. This is due to variability between species, stomatal response to the treatment itself, and to environmental effects. Here we attempt a quantitative evaluation with a defined aerosol application, a paired leaf experimental design, and immobilized stomata.

Attached leaves of poplar were treated with ammonium nitrate aerosol. After 17 or 20 days for deliquescence to develop an aqueous film, leaves were excised and stomata held closed with abscisic acid or open with fusicoccin. Transpiration and stomatal conductance were measured in a greenhouse with a porometer and leaf health was assessed by fluorescence.

Median stomatal conductance was increased significantly, by 60 and 65%, following aerosol loading of 31.3 μg cm^-2^ in ABA- and FC-treated leaves, respectively.

Aerosol induced transpiration, probably associated with a liquid film that lines the stomatal pore and not effectively regulated by stomatal closure, may be significant in magnitude. As aerosol deposition is ubiquitous, and its chemical nature may be changing, this factor should be considered in models of transpiration from leaf to canopy scale.

## Introduction

Unexpected plant mortality in the Anthropocene reflects increasing atmospheric and edaphic drought (Novick et al., 2016; Grossiord et al., 2020). Increased evaporative demand is more important than increasing leaf temperature (T_l_) in driving recent increases in plant mortality (Eamus et al., 2013; Yuan et al., 2019). Leaf to air vapor pressure difference (VPD_l_) represents the driving force for transpiration (E) and is a primary regulator of stomatal aperture (Farquhar, and Raschke, 1978; Grantz, 1990; Lange et al., 1971; Medlyn et al., 2011; Monteith, 1995). The mechanism of stomatal regulation by atmospheric humidity and, particularly, the site of evaporation during stomatal transpiration (Cernusak et al., 2018; Farquhar and Raschke 1978; Grantz 1990; Kaiser 2009; Tyree and Yianoulis, 1980) remain enigmatic despite decades of experimental and analytical research (Aphalo and Jarvis, 1991; Bauer et al., 2013; Buckley, 2005, 2015; Cardoso et al., 2020; Cernusak et al., 2018; Dewar, 2002; Farquhar and Raschke, 1978; Lange et al., 1971; Monteith, 1995; Rockwell et al., 2014; Scoffoni et al., 2017; Tyree and Yianoulis, 1980). Because transpiration affects regional weather, and stomata both respond to weather and control transpiration, quantitative understanding of stomatal water flux is a critical component in understanding climate change (Hetherington and Woodward, 2003; Novick et al., 2016). The continuing uncertainties regarding stomatal responses to atmospheric humidity, and gradients therein, suggest that an additional and previously unrecognized factor may be involved.

Atmospheric aerosols and their deposition to plant canopies are ubiquitous (Grantz et al., 2003; Burkhardt and Grantz, 2017), predate the Anthropocene (Andreae, 2007), and are increasing in flux and hygroscopicity (Hamilton, 2015; Lindberg et al., 1986; Putaud et al., 2010; Tsigaridis et al., 2006). Hygroscopic particles function as cloud and fog condensation nuclei in the atmosphere, and dew condensation nuclei on leaf surfaces, contributing to the thin aqueous film that coats all surfaces (Ewing, 2005; Wylie, 1955; Petters and Kreidenweis, 2007) including leaves, (Burkhardt et al., 2001b; Eiden et al., 1994). Aerosol-induced condensation to foliar surfaces has been visualized using environmental scanning electron microscopy (ESEM; Eiden et al., 1994; Burkhardt and Hunsche, 2013; Grantz et al., 2018).

As deliquescent particles undergo equilibrium transformation to a thin film of saturated solution (Pilinis et al., 1989), the liquid can be seen to undergo ‘salt creep’ (Ivanova and Esenbaey, 2021; Qazi et al., 2019), overcoming the surface tension of hydrophobic leaf surfaces (Kunz, 2010; Pegram and Record, 2007; Roth-Nebelsick, 2007) and entering stomatal pores (Arsic et al., 2020; Burkhardt and Hunsche, 2013; Burkhardt et al., 2012; Burkhardt & Pariyar, 2014 (with videos); Eichert et al., 2008; Park et al., 2023). Diffusion of materials into the stomatal pore along this liquid pathway has been directly observed (Basi et al., 2014; Eichert et al., 1998; Kaiser, 2014). This has the potential to link apoplastic water to the leaf surface, moving the site of evaporation to outside the stomata (Burkhardt, 2010). This bypasses stomatal control and may increase transpiration, particularly as stomata close, because liquid water is more dense than vapor, is incompressible, and does not respond to the changes in stomatal aperture that regulate vapor phase fluxes. This moves the competition between liquid and vapor transport (Rockwell et al., 2014) to within and beyond the stomatal pore, and alters plant water balance (Pariyar et al., 2013; Chi et al., 2022).

Current models of transpiration, from leaf to global scale, explicitly assume that transpiration is dominated by stomatal water vapor flux, and is thus regulated by stomatal aperture (Parlange and Waggoner, 1970). Yet, in the presence of aerosols, higher stomatal conductance (g_s_) per unit stomatal aperture has been observed in two dissimilar genera, *Vicia* and *Sambucus* (Burkhardt et al., 2001; Grantz et al., 2018). In *Vicia*, the heterogeneity of g_s_ (stomatal patchiness) was reduced by aerosol exposure. This patchiness has also been enigmatic (Beyschlag and Eckstein, 1998; Downton et al., 1988; Pospisilova and Santrucek, 1994; Spence, 1987 Weyers and Lawson, 1997) and its apparent reduction by coordination of g_s_ across the leaf surface by hydraulic coupling within the leaf (Buckley and Mott, 2000; Mott et al., 1999; Nardini et al., 2008) and coupling by thin aqueous films across the leaf surface (Grantz et al., 2020), has implications for the regulation of gas exchange (Buckley et al., 1999; Cheeseman, 1991; Cardon et al., 1994; Haefner et al., 1997). These aerosol effects on E and patchiness have been absent from both models of water balance and from calculations associated with porometric and gas exchange measurements.

Effects of aerosol deposition have proven difficult to document over many years of research (Burkhardt et al., 2001; Chi et al., 2022). The aerosol impact cannot be isolated by gas exchange measurements alone, because vapor and liquid flux streams merge at the site of measurement. Identification requires either direct comparison of either flux and aperture, or flux with and without aerosol deposition.

The magnitude of the effect has not been quantified, and may be small. Stomatal responses to the additional increment of aerosol-facilitated transpiration (Δg_s_) may, themselves, confound efforts to identify it (Burkhardt et al., 2001; Pariyar et al., 2013; Chi et al., 2022). For example, robust stomatal closure with increasing VPD_l_ is a defining characteristic of isohydric behavior and Δg_s_ may be sensed similarly to increased VPD_l_. Stomatal response could mask small aerosol effects. Additionally, interspecific differences make generalizations difficult, with beech exhibiting increased sap flow while pine did not (Burkhardt and Pariyar, 2016). During the simultaneous microscopic and gas exchange measurements with *Sambucus nigra* L. (Burkhardt et al., 2001) and with *Vicia faba* L. (Grantz et al., 2018), stomatal responses to VPD_l_ and to aerosol were observed.

In the current study, we have immobilized stomata in either the closed state (with abscisic acid, ABA) or in the open state (with fusicoccin, FC). Within each treatment a broad range of conductances were observed. This provides measurements of the magnitude of Δg_s_ over a broad range of g_s_, and avoids confounding from stomatal response. In this way we quantify the aerosol effect, for the first time, an important first step in incorporating aerosol effects into modeling and calculations associated with porometric instruments.

## Materials and Methods

### Plant material

A hybrid poplar clone (*Populus maximowiczii x nigra*) was grown hydroponically, with solutions renewed weekly. This clone has been identified as relatively isohydric (Xu et al., 2018). Trees were propagated from seedlings in 3 pots with 4 trees per pot (n = 12), and grown in an aerosol-free greenhouse in Bonn, Germany as described previously (Grantz et al., 2018). Trees had been coppiced and grown to about 2 m in height with multiple branches.

A range of leaf sizes was used for experiments (Fig. 1a taken from the distal portion of healthy branches. Adjacent, alternate leaves were selected as pairs on a single branch per tree. Attached leaves, one of each pair, were treated by misting with water or with 25 mM NH_4_NO_3_ solution. This is an effective, and quantifiable, surrogate for aerosol deposition. Misting occurred on 4 September 2023 for the ABA experiment and on 11 September 2023 for the FC experiment. Each leaf being misted was isolated with an aluminum foil cone to avoid spray drift.

**Figure 1.**
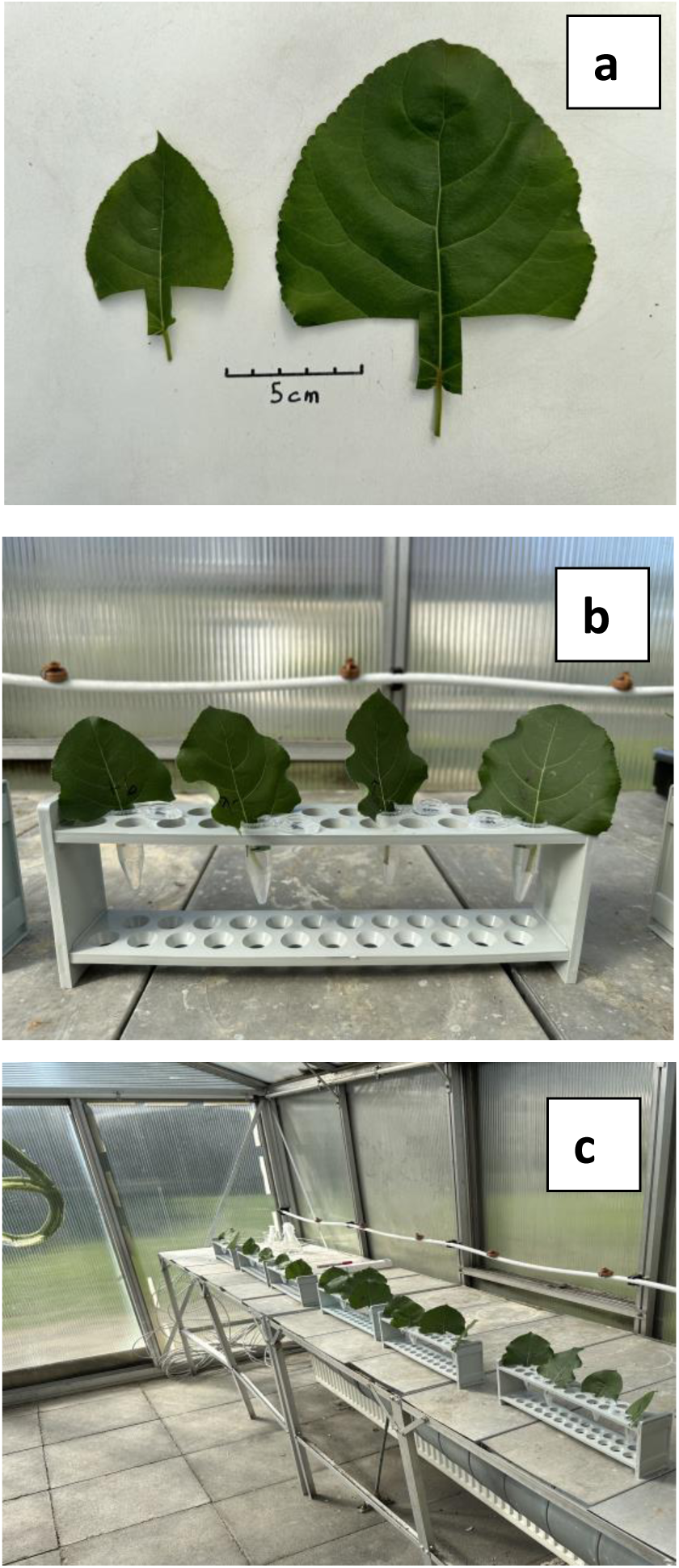
(a) Leaves of all sizes were used for experiments, taken as pairs of adjacent alternate leaves (b) Excised leaves were inserted into Eppendorf tubes; (c) Tubes were held in a row of racks with Control and +Aerosol leaves alternating and measurements conducted sequentially along the rack.

### Porometry and Fluorimetry

A hand held porometer/fluorimeter (Model 600; Licor Inc. Lincoln NE USA) was used for all measurements. Leaf surfaces were allowed to dry prior to the first measurement. The porometer was clamped onto leaves, away from major veins, with the measurement aperture (0.75 cm diameter) fully covered by healthy leaf area on the abaxial surface. No measurements were attempted on the adaxial surface. While transpiration (E) is directly measured with the porometer, we used g_s_ as a measure of water loss as it represents E normalized for leaf to leaf variation in VPD_l_.

Ambient photosynthetic photon fluence rate (PPFR, μmol m^-2^ s^-1^) was measured in the plane of the leaf at the time of each conductance measurement. The porometer was set to the default ‘Auto gsw+F’ in the ‘fast’ preset mode, with stability of 0.005 mol m^-2^ s^-1^ over 2 s. Flow was 150 μmol s^-1^.

For fluorescence measurements, taken concurrently with the conductance measurements, leaf absorptance was set to 0.8 and fraction absorptance of PSII to 0.5. Pulse modulation rate was set to 500 Hz, with flash intensity at 7000. ΦPSII was calculated in the light adapted mode as ΔF/Fm’, and represents the fraction of absorbed PSII photons utilized for photochemistry. Electron transport rate was calculated as the product of PPFR and ΦPSII.

### Measurements

Leaves were excised and measurements obtained on 24 September 2023 for the ABA experiment and on 28 September 2023 for the FC experiment. This resulted in a 20 day exposure to aerosol in the ABA experiment and 17 day exposure in the FC experiment. On measurement days using a new scalpel, leaves were excised from the intact plant at the point of petiole attachment to the stem, quickly submerged in water, petioles recut, and the laminae cut to a spade shape (Fig. 1a). The petiole and leaf tab were inserted into individual Eppendorf tubes held in racks (Fig. 1b). For the ABA experiment, tubes were filled and replenished with a solution of 1 x 10^-5^ M abscisic acid (2-cis,4-trans-Abscisic Acid, 98%; Sigma-Aldrich; St. Louis MO, USA; product 862169). For the FC experiment, tubes were filled and replenished with a solution of 1 x 10^-5^ M fusicoccin (Sigma-Aldrich; Schnelldorf, Germany; product F0537).

Leaves were held in a row of 6 racks arrayed linearly along the north wall of the greenhouse, with two leaf pairs per rack and paired leaves (i.e. Control and +Aerosol) alternating (Fig. 1C). Measurements were conducted sequentially along the row of racks, thus alternating control and treated leaves. Laminae were initially oriented toward the center of the greenhouse, inclined slightly from the vertical, with the adaxial surface uppermost. This orientation was inadvertently partially randomized during repeated attachment of the porometer head.

Temperature (T_l_) and leaf to air vapor pressure deficit (VPD_l_) were recorded at individual leaves during the porometric measurements (Table 1). VPD_l_ increased somewhat during the course of the measurement day along with ambient temperature (not shown), but individual leaf temperature (T_l_) and VPD_l_ were determined primarily by leaf orientation. Light (photosynthetic photon fluence rate, PPFR) on the leaves during measurement was slightly higher in the FC (163 mmol m-2 s-1) than ABA (132 mmol m-2 s-1) experiment (not shown), but VPD_l_ was lower (Table 1) reflecting greater relative humidity on the FC measurement day (not shown). The range of VPD_l_ extended to about 5 kPa in the ABA experiment and to about 3 kPa in FC.

**Table 1.** Comparison of leaf environmental parameters of +Aerosol or Control during combined porometric and fluorimetric measurements of leaves treated with abscisic acid (ABA; upper table) or fusicoccin (FC; lower table) on the day of measurement.

| ABA Experiment | Control<br>N = 216 |  | +Aerosol<br>N = 216 |  | P |
| --- | --- | --- | --- | --- | --- |
|  | Median<sup>1</sup> | 25% -75% | Median | 25% - 75% |  |
| Leaf Temperature<br>T<sub>l</sub> (°C) | 27.96 | 24.94 –<br>29.71 | 28.02 | 24.88 –<br>29.66 | ns |
| Vapor Pressure<br>Deficit, VPD<sub>l</sub><br>(kPa) | 2.33 | 1.96 – 2.94 | 2.37 | 1.95 – 2.92 | ns |
| FC Experiment | Control<br>N = 72 |  | +Aerosol<br>N = 72 |  |  |
|  | Mean<sup>2</sup> | Std. Dev. | Mean | Std. Dev. |  |
| Leaf Temperature<br>T<sub>l</sub> (°C) | 27.520 | 2.616 | 27.246 | 2.579 | 0.0038 |
| Vapor Pressure<br>Deficit, VPD<sub>l</sub><br>(kPa) | 1.817 | 0.532 | 1.737 | 0.513 | 0.0030 |

<sup>1</sup>These data failed the Shapiro-Wilk Test for normality (P<0.05). Median separation (P<0.05) was by Wilcoxon Signed Rank Test.
<sup>2</sup>Mean separation (P<0.05) was by paired, two tailed t Test.

In the ABA experiment there was no significant difference in T_l_ or VPD_l_ between +Aerosol and Control leaves (Table 1). At higher VPD_l_ the +Aerosol leaves trended towards greater VPD_l_ (Fig. 2A). In the FC experiment, T_l_ and VPD_l_ were significantly lower in the +Aerosol treatment (Table 1). Mean T_l_ was 0.27°C cooler, resulting in mean VPD_l_ 0.08 kPa lower in +Aerosol leaves. This was observed at all levels of VPD_l_ (Fig. 2B) and reflects aerosol-induced transpiration, which was greater in FC than in ABA.

**Figure 2.**
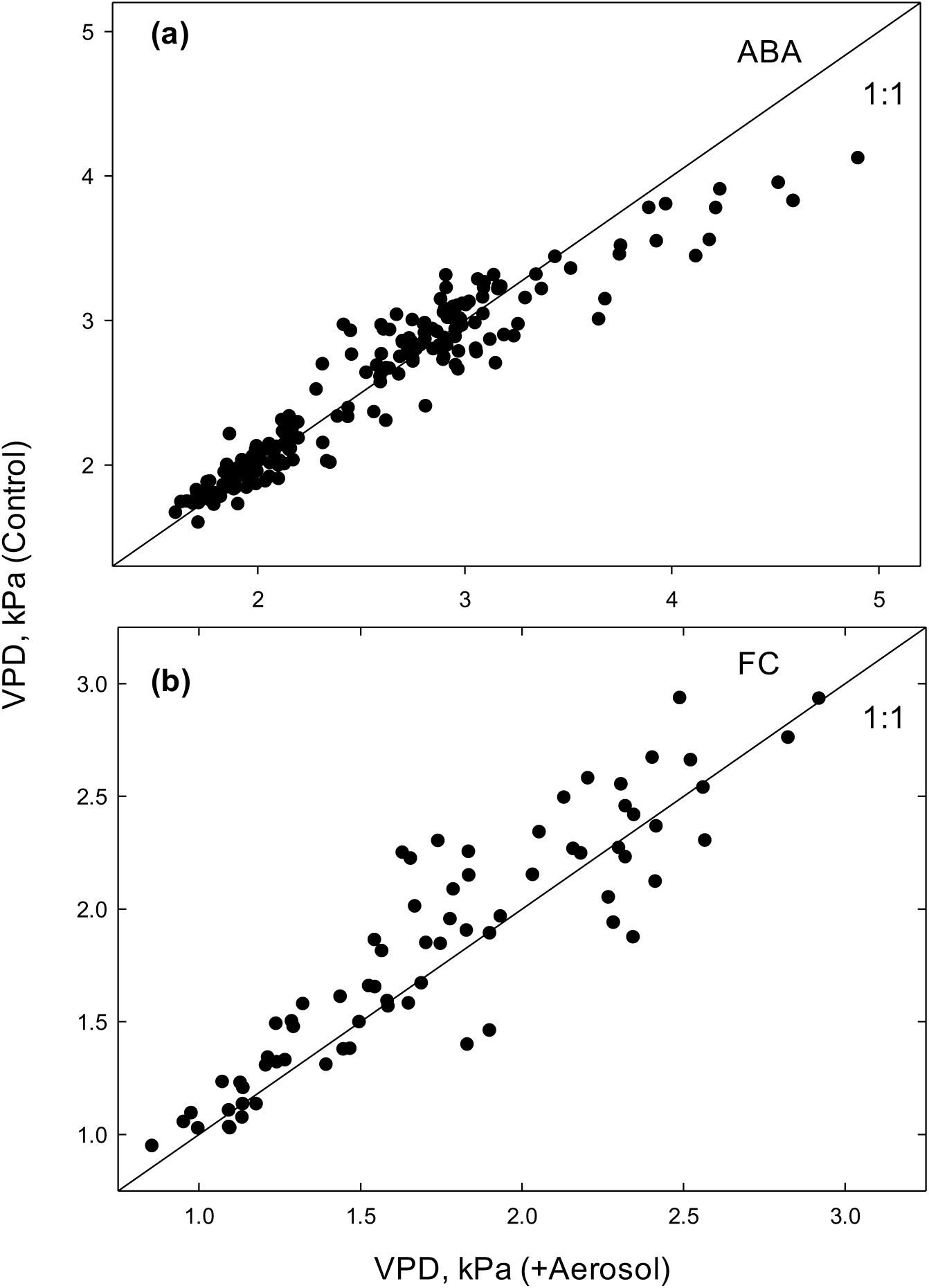
Relationship between VPD for control leaves (Y axis) and VPD for aerosol treated leaves (X axis) in the abscisic acid (ABA) experiment (a) and the fusicoccin (FC) experiment (b).

### Solutions

Both ABA and FC were stored at −20°C in dry form and following solubilization to 1 x 10^-2^ M in solvent. These stock solutions were diluted to final concentration the night before measurements. Final solutions contained 0.1% Methanol (ABA) and 0.1% Ethanol (FC).

### Data analysis

The ABA experiment and the FC experiment were conducted and analyzed independently. For each, a sample size of 12 plants per aerosol treatment was utilized (n=12). A paired sample protocol was used for data analysis (paired t-test; Sigma Plot v. 12.5). The Shapiro–Wilk test was used to test data for normality. For normally distributed data, mean separation was evaluated by two-tailed Student’s t-test. For non-normally distributed data, median separation was evaluated nonparametrically using the Wilcoxon signed rank test.

Figures were prepared using Sigma Plot v. 12.5. The histograms display different bar ranges, in order to preserve 10 bins for all treatments while encompassing all data. No outliers were excluded at any stage of analysis.

## Results

### Effect of uptake of ABA or FC on excised leaves

Pair-wise comparison of immobilized stomata of leaves treated with water or with an aerosol mist of NH_4_NO_3_ allowed separation of relatively subtle differences in transpiration (E) and apparent stomatal conductance (g_s_). These aerosol effects have previously been difficult to resolve.

Treatment with ABA reduced median E by 87.0% and 88.3%, relative to FC (control and +Aerosol leaves, respectively; cf. Tables 2). Treatment with ABA reduced median g_s_ by 91.4 and 91.7%, relative to FC (control and +Aerosol leaves, respectively; cf. Tables 2). In closed stomata taking up ABA, Δg_s_ was 0.0075 mol m^-2^ s^-1^ and in open stomata taking up FC, Δg_s_ was 0.095 mol m^-2^ s^-1^.

### Effect of aerosol on E

In closed stomata (ABA experiment; Table 2), measured median levels of E were significantly (30.0%) greater in +Aerosol leaves than in Control leaves (P = 0.002). In the parallel experiment with open stomata (FC experiment; Table 2), median E in +Aerosol was 37% greater (P < 0.001). Because VPD_l_ is a determinant of E, and varied non-systematically among leaves (see Materials and Methods), further analysis was conducted on the basis of apparent stomatal conductance (g_s_), which normalized for individual leaf VPD.

**Table 2.** Comparison of observations of +Aerosol or Control during combined porometric and fluorimetric measurements of leaves treated with abscisic acid (ABA; upper table) or fusicoccin (FC; lower table) on the day of measurement.

| **ABA Experiment** | **Control**<br>**N = 216** |  | **+Aerosol**<br>**N = 216** |  | **P** |
| --- | --- | --- | --- | --- | --- |
|  | **Median<sup>1</sup>** | **25% -75%** | **Median** | **25% - 75%** |  |
| Stomatal Conductance, $g_s$ (mol m<sup>-2</sup> s<sup>-1</sup>) | 0.0125 | 0.0055 - 0.0318 | 0.0200 | 0.0075 - 0.0398 | 0.002 |
| Transpiration, E (mmol m<sup>-2</sup> s<sup>-1</sup>) | 0.322 | 0.131 – 0.791 | 0.460 | 0.165 – 0.940 | 0.002 |
| $\Phi_{psII}$ | 0.683 | 0.630 – 0.740 | 0.702 | 0.622 – 0.736 | ns |
| Electron Transport Rate, ETR; (μmol m<sup>-2</sup> s<sup>-1</sup>) | 35.32 | 13.39 – 54.15 | 36.24 | 11.87 – 57.37 | ns |
| **FC Experiment** | **Control**<br>**N = 72** |  | **+Aerosol**<br>**N = 72** |  | **P** |
|  | **Median<sup>1</sup>** | **25% -75%** | **Median** | **25% - 75%** |  |
| Stomatal Conductance, $g_s$ (mol m<sup>-2</sup> s<sup>-1</sup>) | 0.145 | 0.0317 – 0.486 | 0.240 | 0.081 – 0.540 | <0.001 |
| Transpiration, E (mmol m<sup>-2</sup> s<sup>-1</sup>) | 2.473 | 0.739 – 5.133 | 3.924 | 1.524 – 5.621 | <0.001 |
| $\Phi_{psII}$ | 0.694 | 0.659 – 0.706 | 0.692 | 0.660 – 0.704 | ns |
| Electron Transport Rate, ETR; (μmol m<sup>-2</sup> s<sup>-1</sup>) | 41.926 | 31.902 – 59.553 | 43.978 | 30.429 – 63.583 | ns |

<sup>1</sup>These data failed the Shapiro-Wilk Test for normality (P<0.05). Median separation (P<0.05) was by Wilcoxon Signed Rank Test.

### Effect of aerosol on g_s_

In closed stomata (ABA experiment; Table 2), measured median levels of g_s_ were significantly (60.0%) greater in aerosol-treated (+Aerosol) leaves than in leaves misted with water (Control; P = 0.002). The median increase in g_s_ (Δg_s_) was 0.0075 mol m^-2^ s^-1^. In the parallel experiment conducted with FC (Table 2), median g_s_ was increased in +Aerosol by 65.5% (P < 0.001). The median Δg_s_ was 0.095 mol m^-2^ s^-1^.

In the ABA experiment, the greater g_s_ in +aerosol treatment was observed at all levels of g_s_, but particularly in the middle range (0.01-0.05 mol m^-2^ s^-1^; Fig. 3a). 61% of observations in +Aerosol lay between 0.011 – 0.078 mol m^-2^ s^-1^, while 41.0% of Control observations lay in the lightly smaller range of 0.016 – 0.078 mol m^-2^ s^-1^ (cf. Fig. 4ab).

**Figure 3.**
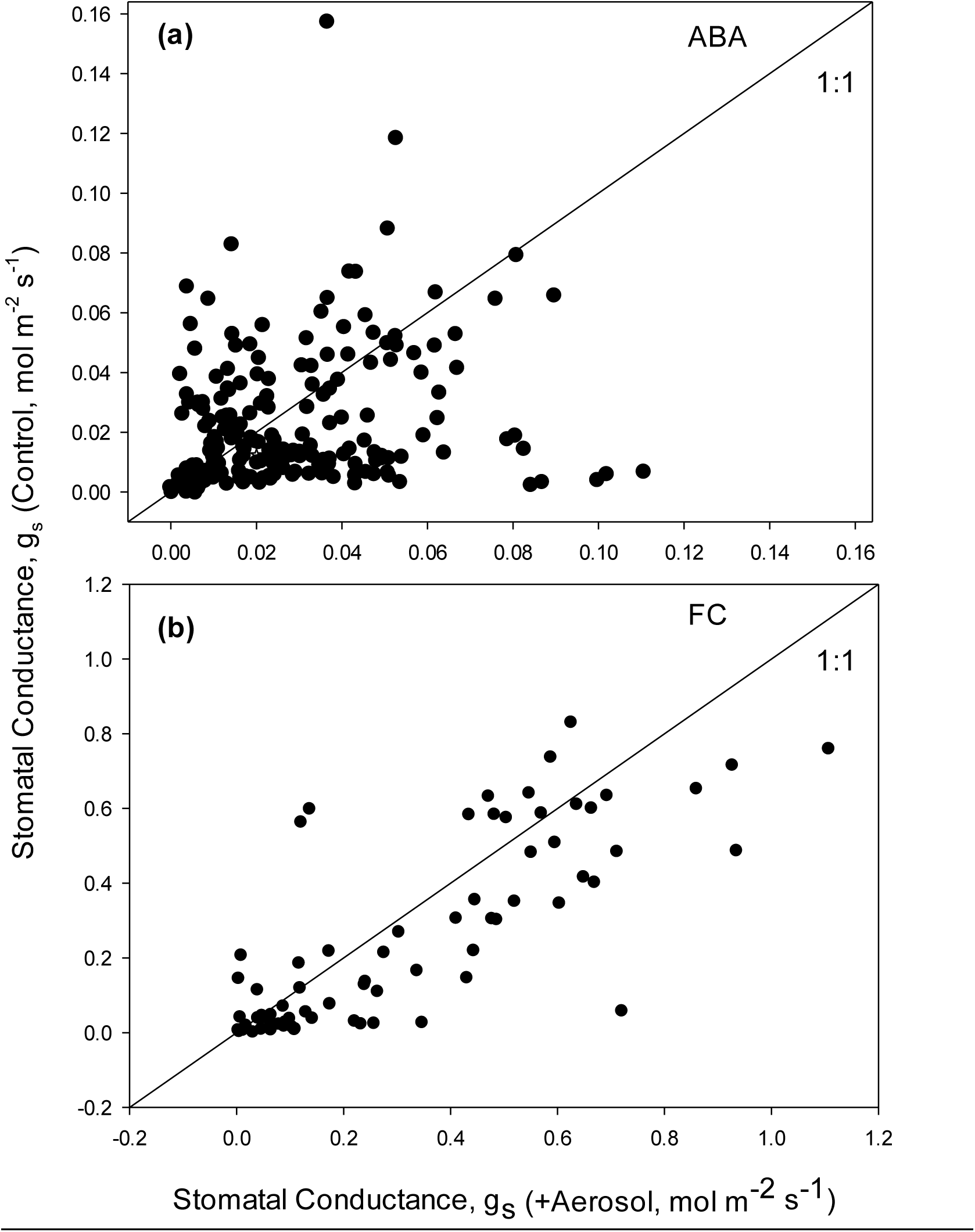
Relationship between stomatal conductance of control leaves (Y axis) and aerosol treated leaves (X axis) in leaves (a) closed with abscisic acid (ABA) and leaves (b) opened with fusicoccin (FC).

**Figure 4.**
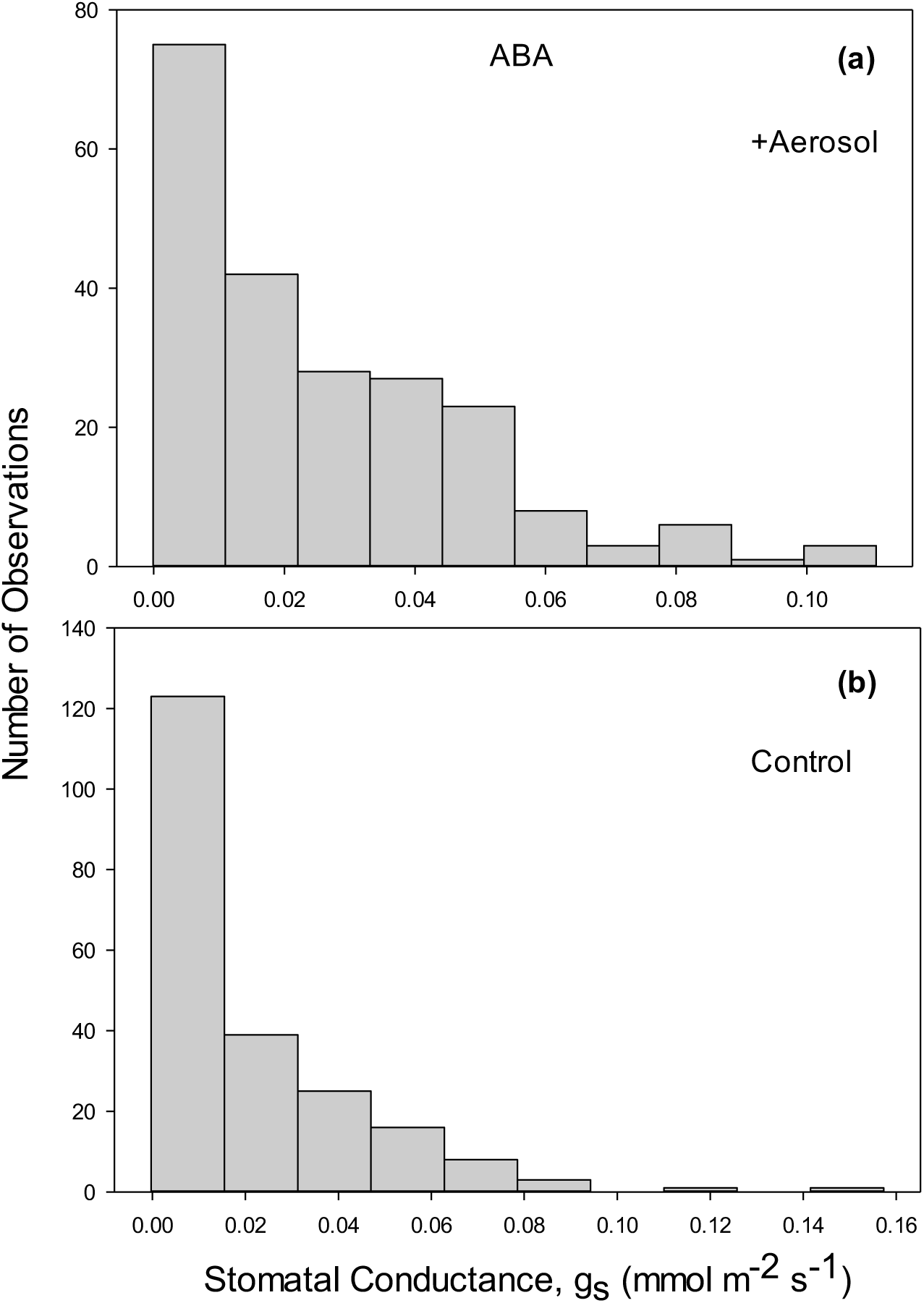
Distribution of stomatal conductances of paired leaves closed with abscisic acid (ABA), following exposure to (a) aerosol or (b) water.

In the FC experiment, the skewing of g_s_ towards larger values in the +aerosol leaves was more evident at all levels of g_s_ (Fig. 3B). 52.8% of +Aerosol observations lay between 0.155 – 0.707 mol m^-2^ s^-1^ (Fig. 5a) and 43.0% in the slightly smaller range of 0.156 – 0.695 mol m^-2^ s^-1^ in the Control (Fig. 5b).

**Figure 5.**
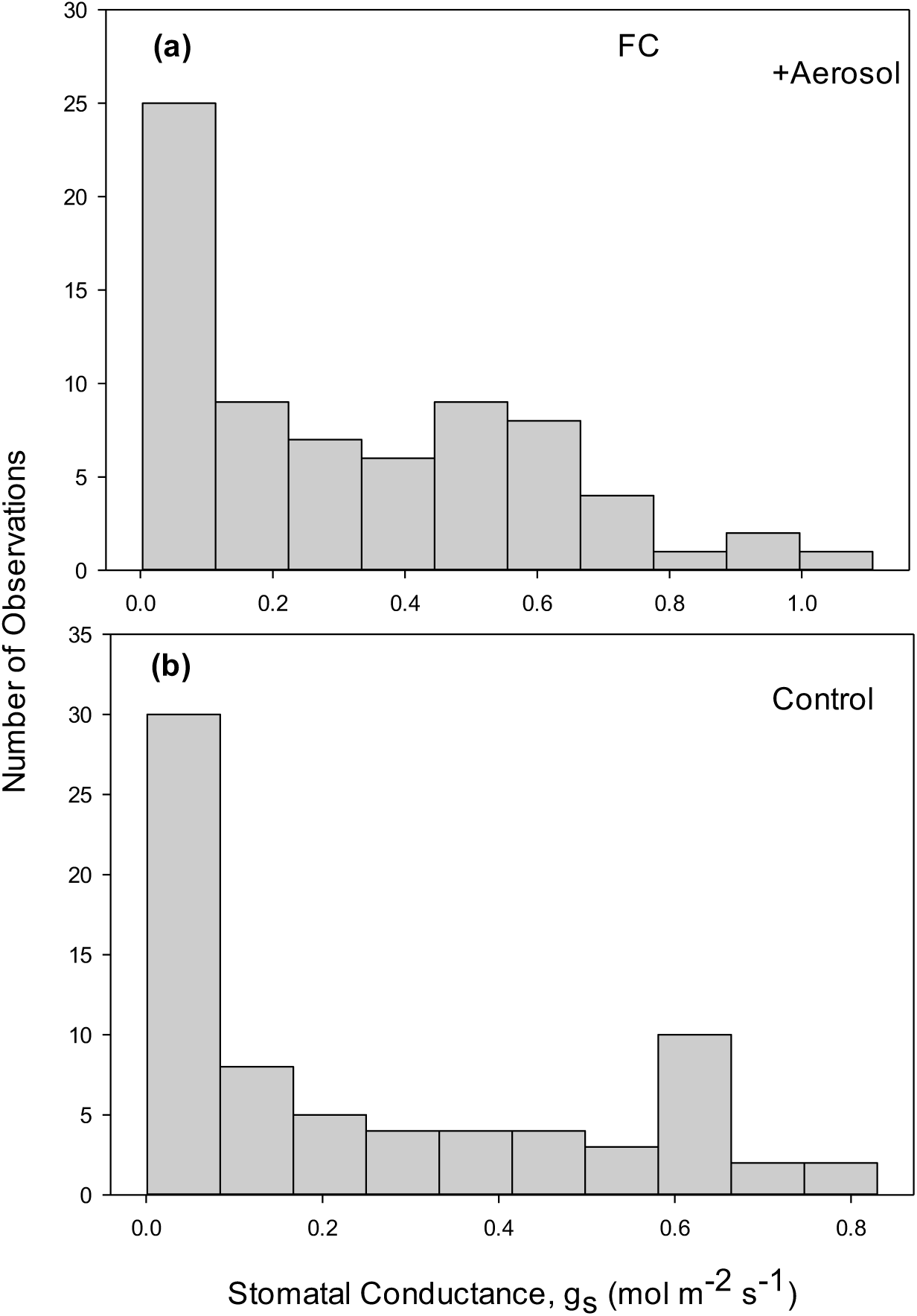
Distribution of stomatal conductances of paired leaves opened with fusicoccin (FC), following exposure to (A) aerosol or (B) water.

### Photochemical health of the leaves

The capability of the LI600 to measure Chlorophyll ***a*** fluorescence simultaneously with porometry was utilized as a check on the health of the excised leaves throughout the day. There were no differences in ΦPSII or ETR between Control and +Aerosol leaves in either the ABA or FC experiment. While ΦPSII was similar on the ABA and FC measurement days, ETR was greater in FC due to the greater PPFR at the leaf surfaces (Table 2).

In the ABA experiment, there was no relationship between VPD_l_ and g_s_ in either +Aerosol (Fig. 6a) or Control leaves (Fig. 6b).The range of VPD_l_ extended to >4 kPa. In the FC experiment (Fig. 7), apparent stomatal conductance (g_s_) declined asymptotically to the origin with increasing VPD_l_.

**Figure 6.**
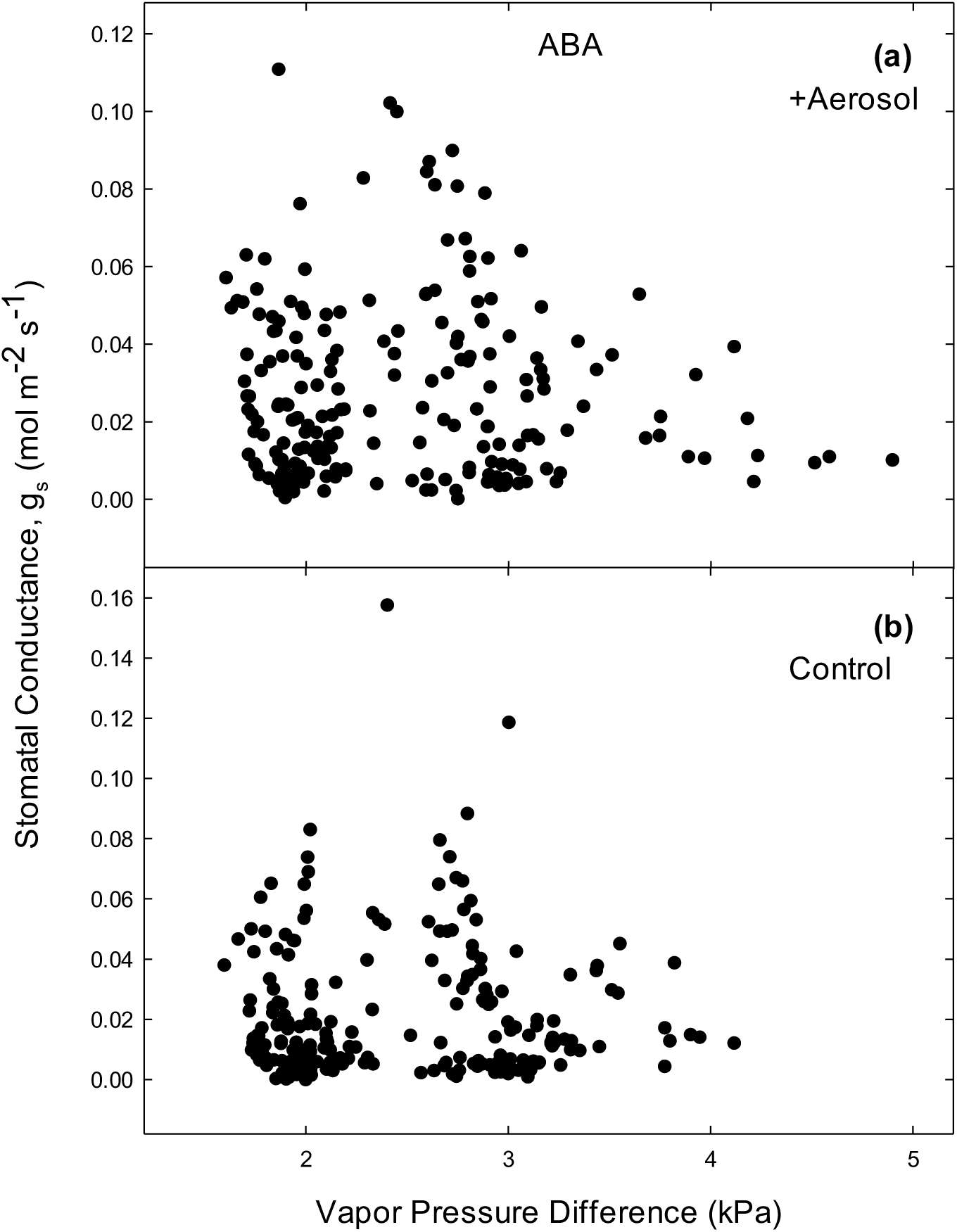
Relationship between stomatal conductance of (a) aerosol treated leaves or (b) control leaves and vapor pressure deficit (VPD) in leaves closed with abscisic acid (ABA).

**Figure 7.**
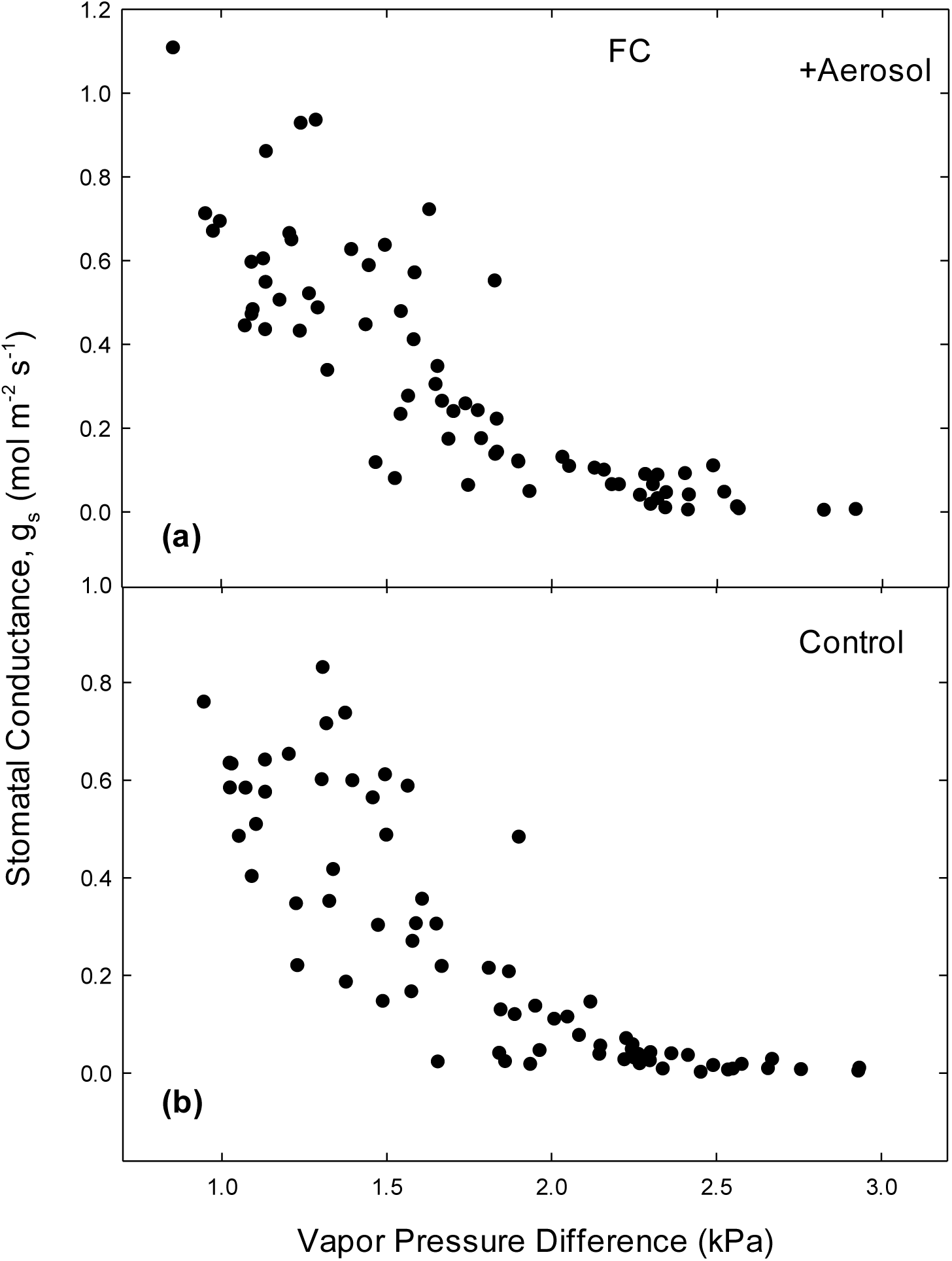
Relationship between stomatal conductance of (a) aerosol treated leaves or (b) control leaves and vapor pressure deficit (VPD) in leaves opened with fusicoccin (FC).

The aerosol-induced increment of stomatal conductance, Δg_s_ (+Aerosol minus Control value calculated for leaf pairs) declined slightly but not significantly with increasing VPD_l_ in both ABA and FC experiments (Fig. 8).

**Figure 8.**
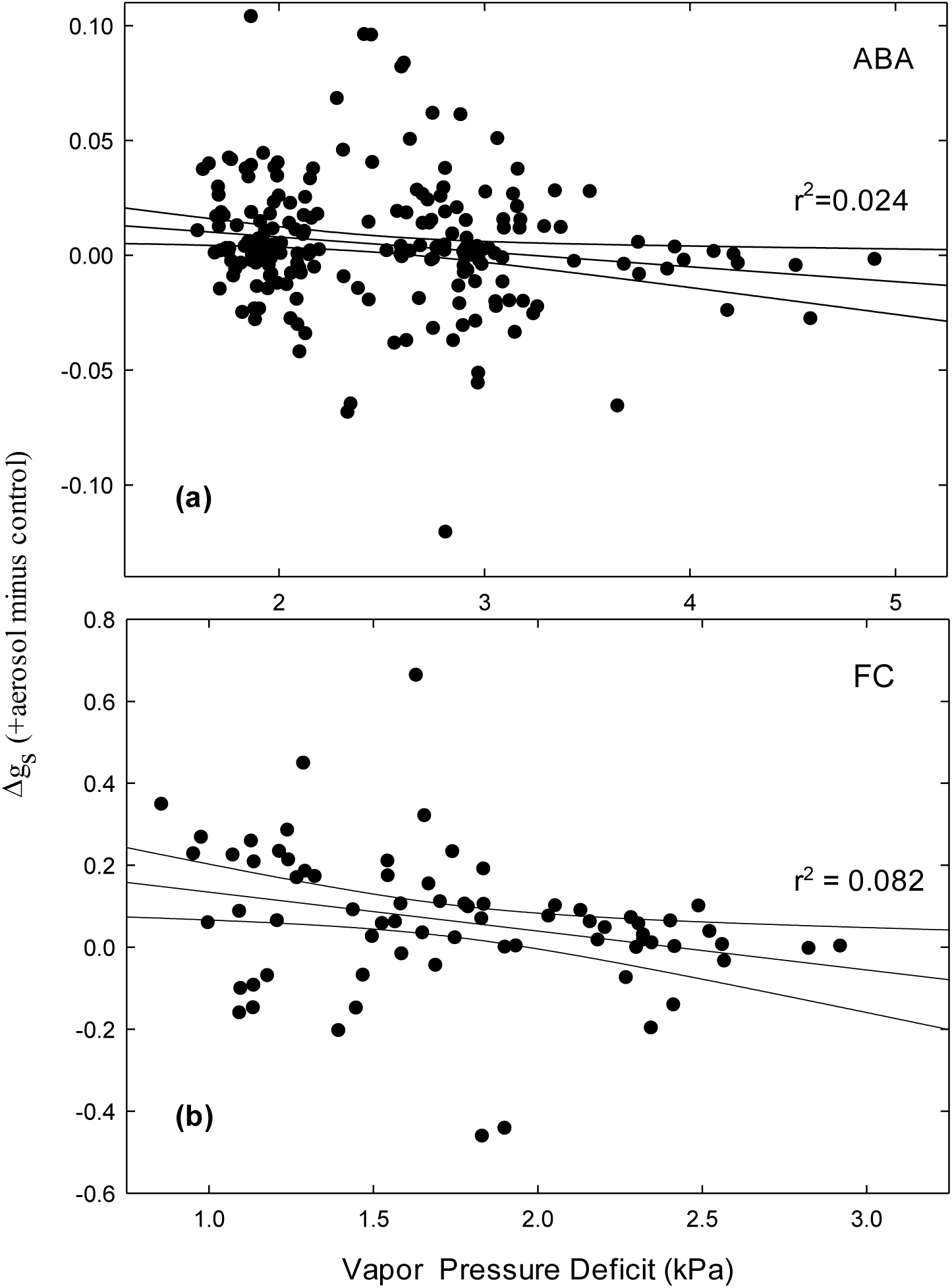
Relationship between the aerosol impact on gs and Vapor Pressure Deficit (kPa) in leaves closed with ABA or opened with FC.

## Discussion

Our large number of biological replicates (12 trees) and measurements on paired leaves (n=216 in the ABA experiment and n=72 in the FC experiment), and the use of immobilized stomata, allowed us to quantify the aerosol impact on transpiration (ΔE) and stomatal conductance (Δg_s_). Although E is the directly measured parameter, our analysis is based on apparent stomatal conductance (g_s_) to reduce confounding by differences in individual VPD_l_. Aerosol effects on E were just as significant (P = 0.002 and P < 0.001) as effects on g_s_ (also P = 0.002 and P < 0.001), in ABA and FC experiments, respectively. However, the magnitude of effects on E was smaller, 30% and 37%, vs. 60% and 66% for g_s_.

In both ABA and FC experiments, g_s_ was increased by aerosol over all levels of g_s_, but particularly in the middle range, 61% vs. 41% of observations in ABA and 53% vs. 43% in FC. Very large values of g_s_ were also more prevalent in the +Aerosol treatment, increased from 2% to 5% in the ABA experiment and from 6% to 11% in the FC experiment.

The ABA experiment is similar to studies of minimum leaf conductance conducted with detached leaves drying in the laboratory. Under those conditions, it has been consistently observed that minimum conductance increases following aerosol exposure (Burkhardt & Pariyar, 2014; Burkhardt *et al*., 2018; Grantz *et al*., 2018; Chi *et al*., 2022). In the current experiment, median Δg_s_ in the ABA experiment was positive in the presence of aerosol. This is likely to represent additional liquid water flux through the stomata. Cuticular conductance is typically <0.001 mol m^-2^ s^-1^ (Duursma et al. 2018) and is not likely to affect these results, although aerosol impacts on cuticular permeability have not been rigorously evaluated. Nevertheless, the cuticle is complex and under-studied (Fernandez et al., 2017; Kerstiens 1996) and cuticular conductance is known to vary with altitude (Anfodillo et al., 2002) and turgor (Boyer 2015b).

The excised leaves remained viable throughout the measurements. The levels of g_s_ observed in the FC experiment were similar to those observed in intact leaves of these trees (Grantz, unpublished observations. g_s_ did not decline and measures of photochemical competences remained steady over the course of the measurement day. These measures, ΦPSII and ETR, did not differ between Control and +Aerosol leaves in ABA or FC. The ABA treated leaves were clearly immobilized in the closed state, with considerable variability but no relationship with VPD_l_. This variability could be due in part to variable application of the aerosol mist. It is less certain that the FC treated leaves were completely immobilized in the open state. While there was considerable variability in g_s_ including some nearly closed pores, overall g_s_ in the FC experiment exhibited a hyperbolic decline to the origin with increasing VPD_l_. This is similar to the relationship commonly observed in intact and untreated leaves (Aphalo and Jarvis, 1991; Grantz, 1990; Monteith, 1995). Despite the range of g_s_, in both ABA and FC experiments, similar biological properties of the paired leaves overcame this variability to reveal the aerosol effects.

These data support previous conclusions that aerosol deposition increases water flux through the stomata (e.g. Burkhardt and Grantz, 2017). The decline of Δgs with VPD_l_ in both ABA and FC may reflect the multiple impacts of atmospheric humidity (Grantz, 1990; Monteith, 1995) and has been consistently observed (Burkhardt et al., 2001; Pariyar et al., 2013; Grantz et al., 2018). Evaporative demand increases with VPD_l_ while the evaporating surface wetted by the deliquescing aerosol contracts with concomitantly declining relative humidity (Burkhardt et al., 2023). Thus the magnitude of the aerosol effect may depend on many factors, including stomatal morphology, environmental conditions, and aerosol loading. Here we have applied about 31.3 μg cm^-2^ of a highly hygroscopic aerosol, based on leaf surface water holding capacity at runoff for these poplar leaves (Chi, unpublished results of 15.6 ± 0.6 μl cm^-2^; n = 20) and the concentration of NH_4_NO_3_ in the misting solution.

This solution dried quickly with the aerosol remaining on the attached leaves for nearly two weeks prior to measurement. Under these conditions we found that apparent stomatal conductance increased by about 60-65% in both open and closed stomata. The magnitude of additional conductance (Δg_s_) was proportional to baseline conductance (g_s_) as it varied with the ABA or FC treatment. Median Δg_s_ was reduced by 92.1% between FC and ABA, while gs was reduced by 91.7% and 91.4% (in +aerosol and Control leaves, respectively). This proportionality remains somewhat unexplained, as g_s_ is suggested to have less effect on the aerosol mediated liquid pathway than on the traditional vapor phase pathway. Nevertheless, this provides the first quantitative assessment of the aerosol effect on E and g_s_ in the absence of stomatal response to the stimulus being measured.

We conclude that aerosol deposition sets up a component of transpiration that is ubiquitous in nature but underrepresented in clean air laboratory experiments. This is carried by a thin liquid film, estimated to be on the order of 1 x 10^-7^ m thick (Burkhardt, 2010), that lines the throat of the stomatal pore, providing a liquid pathway from mesophyll to leaf surface. Direct evaporation thus occurs directly into the leaf boundary layer rather than into the substomatal space, altering the competition between vapor and liquid phase fluxes (Rockwell et al., 2014) and substantially reducing stomatal control of E. Incorporation of such concepts into models of water balance at all scales is urgently needed.

## Acknowledgments

DAG was supported by the Deutsche Forschungsgemeinschaft (DFG, German Research Foundation), grant number: 446535617 and unrestricted research funds provided by the University of California at Riverside. C-JEC was funded by Deutscher Akademischer Austauschdienst (DAAD, German Academic Exchange Service), grant number: 57440921. JB was funded by the Deutsche Forschungsgemeinschaft (DFG, German Research Foundation), grant number: 446535617.

## Competing interests

We declare no conflict of interests.

## Author contributions

All authors developed the research design and methods. CEC developed the plant system. JB and DAG conducted the experiments. DAG performed data analysis and wrote the paper with inputs from all authors. All authors have read and agreed to the submitted version of the manuscript.

## Notes

### Competing Interest Statement

The authors have declared no competing interest.

